# Measuring the effects of age on contrast suppression

**DOI:** 10.1101/2024.08.16.608357

**Authors:** Martin T. W. Scott, Alex R. Wade, Heidi A. Baseler

## Abstract

The apparent contrast of a suprathreshold central stimulus can be reduced by a surrounding stimulus - a phenomenon known as surround suppression. When stimuli are presented at the fovea, this effect reportedly increases in strength with age. The underlying mechanism proposed for this age dependence, a change in the balance of inhibition and excitation in cortex, makes this phenomenon potentially interesting as a biomarker of neurological dysfunction. Here, we attempt to repeat these measurements, but we use stimuli designed to control for untuned overlay masking, a different form of suppression thought to be of pre-cortical origin. We measured contrast matching thresholds in twenty younger (*<* 30) and seventeen older (*>*60) observers. Across all observers, we find weak suppression that has little or no orientation tuning and, importantly, no dependence on age. Our findings contradict those from earlier studies and suggest that effects relating to age may be dependent on the temporal parameters of this stimulus, and could arise from effects on other pre-cortical mechanisms.

## Introduction

Currently, the global average human life expectancy is 71 years (The World Bank, 2022), and it is projected to be beyond 80 years for newborns by 2050 in the USA (Olshansky et al., 2009) and UK (Office for National Statistics, 2019). This trend calls for an improved understanding of the unique perceptual challenges faced by the elderly. In visual perception, age-related changes in visual acuity and contrast sensitivity are mostly driven by changes to the optical media of the eye (Owsley, 2011). However, there is clear evidence that human visual cortex is subject to structural and functional senescence (Brewer & Barton, 2012), meaning that investigations of post-retinal visual processing across the lifespan may help us to understand how to prevent, treat, or palliate age-related changes in visual perception. One line of enquiry is the perceptual normalisation processes that rely on the delicate balance between neuronal excitation and inhibition, as this balance is known to be disrupted in ageing primates (Fu et al., 2010; Schmolesky et al., 2000). In the context of luminance contrast, “normalisation” refers to the division of a neuron’s sensory inputs by the inputs received from a larger pool of inhibitory neurons. This acts to adjust neural sensitivity to the prevailing visual scene, preserving sensitivity over a wide range of possible visual inputs. It has been suggested that this form of normalisation is a canonical computation – a fundamental neural operation that is enacted throughout the brain (Carandini & Heeger, 2011). If these mechanisms desensitise or diminish with age, it could produce age-related changes in visual perception and other sensory modalities, but also in memory and cognition. Therefore, measuring the action of these mechanisms using relatively simple visual experiments can inform our understanding of the general effect of age on the human brain.

One example of neuronal inhibition in humans is surround-suppression of contrast. When viewing a spatial texture surrounded by a texture of greater contrast, the apparent contrast of the central texture is reduced (or “masked”) (Cannon & Fullenkamp, 1991; Chubb et al., 1989; Petrov & McKee, 2006; Xing & Heeger, 2001). This effect *surround suppression* appears to be physiologically distinct from overlay masking, a form of suppression produced when two stimuli are overlaid (Petrov et al., 2005). Unlike overlay masking, surround suppression operates over long ranges (neighbouring receptive fields) and is tuned for orientation and spatial frequency. This tuning implies a role of elongated receptive fields which are found no earlier than the primary visual cortex. In comparison, overlay masking operates within overlapping receptive fields and is spatially untuned, meaning it could arise as early as the centre-surround receptive fields of the bipolar and retinal ganglion cells. When measuring contrast detection thresholds, many report that surround suppression is weak or effectively absent in the fovea (Meese & Hess, 2004; Petrov et al., 2005; Saarela & Herzog, 2008), but it increases in strength into and beyond the perifovea (Petrov et al., 2005; Snowden & Hammett, 1998). However, there is very little work investigating the tuning of surround-suppression of contrast supra-threshold at the fovea.

There have been several reports suggesting that surround suppression strengthens with age. Initially reported by Karas and McKendrick (2009) using noise stimuli, this pattern of results has also been replicated using drifting luminance gratings (Karas & McKendrick, 2012), and static gratings (Karas & McKendrick, 2011), with the latter work finding the age effect to be present even when the centre and surround gratings are 180° out of spatial phase. Probing a subset of spatiotemporal tuning characteristics, Karas and McKendrick (2015) reported that older adults experience enhanced suppression primarily at low contrasts (centres of 20% contrast), and that surrounds at the same or twice the contrast produce the greatest age-related difference in suppression strength. The same study also found the age difference in surround suppression to be most severe with shorter stimulus presentation times of 150ms, but still observable at a duration of 500ms. Although, later work has indicated that very short presentation times (*∼*40ms) abolish the age-related difference (Pitchaimuthu et al., 2017). Overall, this body of work has often found that surround suppression strengthens with age, opposite to evidence suggesting a weakening of spatial inhibition in older adults in the domain of motion perception(Betts et al., 2005). There is also some evidence from macaque electrophysiology that the orientation tuning of surround suppression broadens with age (Fu et al., 2010). If this is the case, we would simultaneously expect surround suppression to increase in strength but become less orientation specific in older adults. To our knowledge, only Nguyen et al. (2020) has investigated this possibility, but they were unable to replicate the overall effect of age on suppression strength, and did not find any difference in the orientation tuning of surround suppression.

Importantly, when Petrov et al. (2005) measured the distinct spatial tuning properties of surround suppression and overlay masking, they designed their stimuli to reduce cross-contamination between these two suppression types. Primarily, they introduced a gap between the central ‘probe’ region and the surround (equal to one wavelength of the grading stimulus) to reduce the effect of overlay masking when using surround stimuli. This manipulation avoids any confounding effect of overlay masking by reducing the influence of neuronal receptive fields that cover both the centre and the surround. It also prevents complications relating to the perceptual segmentation of the centre from the surround (Appelbaum et al., 2008). They also included a persistent cue to the spatial location of the target which removed spatial uncertainty for low-contrast trials (Petrov et al., 2006). By removing these confounds, the authors effectively isolated and characterised two distinct mechanisms of contrast gain control: an initial, spatially untuned mechanism called ‘overlay masking’ and a later, spatially tuned surround suppression mechanism. It is possible that the absence of these controls in stimuli used by other groups (principally the lack of a one cycle gap) may explain some of the paradoxical effects that have been observed where centers and surrounds directly/closely abut each other.

Using stimuli designed to preclude contributions from overlay masking and spatial uncertainty, the present experiment seeks to characterise the orientation tuning of supra-threshold surround suppression at the fovea, and test the effect of age on the strength and tuning of surround suppression. Using a contrast matching experiment, we psychophysically assessed the strength of surround suppression in younger and older observers using both orthogonal and collinear surrounds. Our stimuli were similar to those used in previous related experiments: luminance contrast gratings centred on the fovea, but with the addition of a mean luminance gap spanning 1-cycle (1*λ*) of the grating’s spatial frequency to prevent overlay masking. If surround suppression alone is the main driver of the age related changes to contrast suppression at fixation, then we expected to find a significant difference in suppression strength between age groups despite this manipulation.

## Methods

### Participants

Seventeen older observers (mean: 69, range: 60 – 81) and twenty younger observers (mean: 18.8, range: 18 – 22) with self-reported normal or corrected-to-normal visual acuity and no personal history of neurological disease or disorder were recruited. This age-range was selected to mirror that of previous studies that have found an effect of age on surround suppression (Karas & McKendrick, 2011, 2015). Ethical approval for the study was given by the Department of Psychology at the University of York, and all observers were reimbursed for their time and travel costs.

### Stimuli and apparatus

The spatial parameters of the stimuli used are illustrated in Figure 1. Stimuli consisted of centrally presented achromatic luminance gratings and annuli with a spatial frequency of 3.33 cycles per degree (cpd). Central gratings always subtended 0.6 degrees of visual angle such that they were precisely 2 cycles of luminance modulation. Central gratings were surrounded by a thin (2 pixel) grey ring to prevent uncertainty pertaining to the location of the target stimulus, which is known to bias detection thresholds (Pelli, 1985; Petrov et al., 2006). When presented, the surrounding annulus subtended 3.2 degrees of visual angle. The outer and inner edges of all stimuli were smoothed by a raised-cosine mask, and the inner and outer edges of the raised cosine plateau were separated by 0.3 degrees of mean luminance (*∼*1*λ*). Stimuli were displayed on a ViewPixx 3D Lite display (1920×1080, 120Hz) (https://vpixx.com/products/viewpixx-3d/), via an Apple Mac Pro 6.1 running macOS High Sierra (version 10.13.6). The display used 8-bits of grayscale resolution and was gamma corrected using a Minolta LS110 photometer. The mean luminance of the display after correction was approximately 50cd/cm^2^. All stimuli were created and presented using the Python programming language (https://www.python.org/) and PsychoPy3 (https://www.psychopy.org/). Observers were seated 1 metre away from the display and used a chin rest while the experiment was in progress. All stimuli were viewed binocularly. Despite wearing correction, two observers in the older age group had difficulty seeing the thin grey ring at a distance of one metre, and were moved closer to the display (*∼*0.68m), but all grating stimuli were temporarily rescaled to accommodate this.

**Figure 1:**
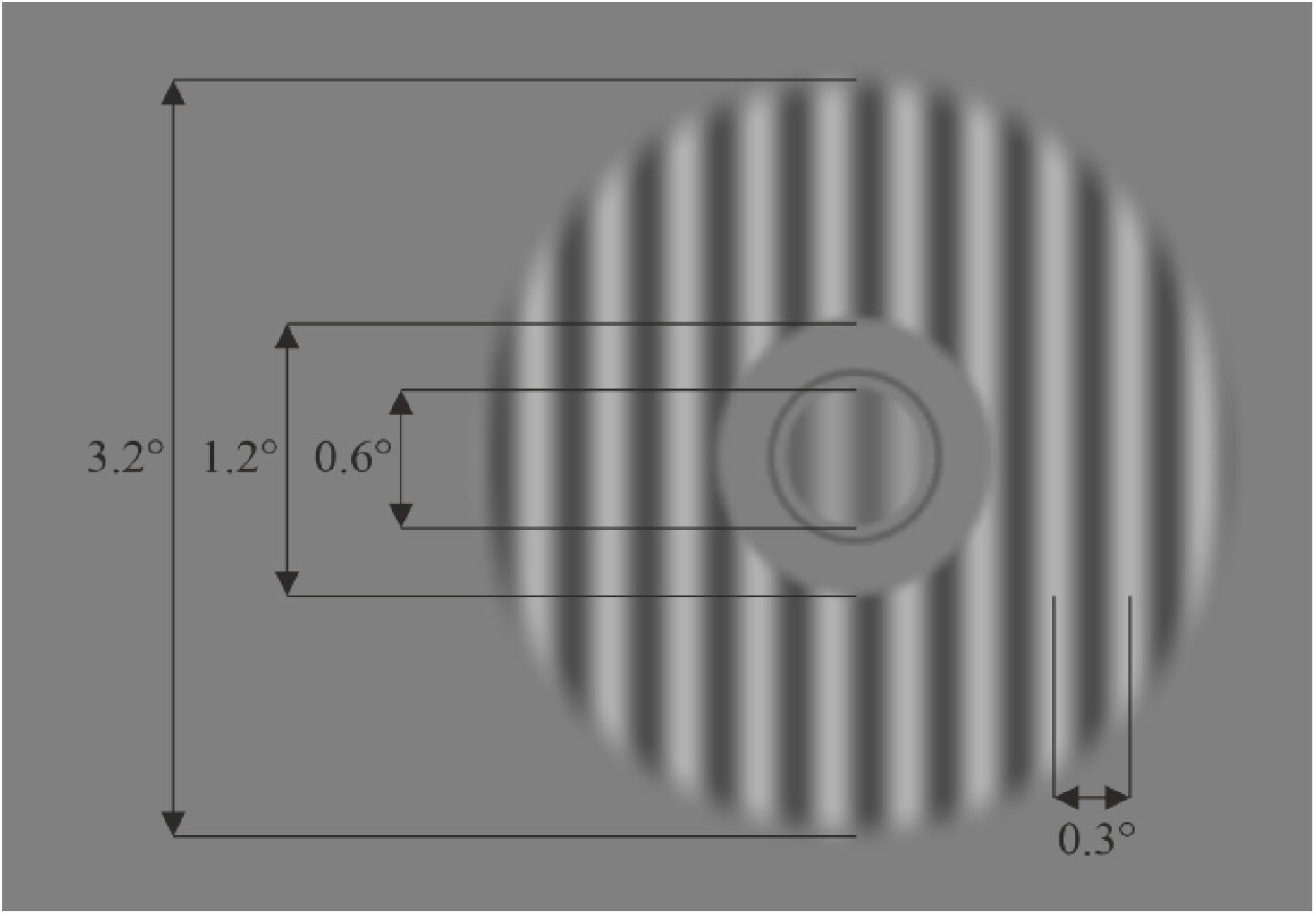
Layout of the stimulus used for our contrast matching experiment. The same central grating and thin ring were used when measuring contrast detection thresholds.

### Procedure

First, to ensure all observers would be able to perceive the stimuli used in the contrast matching task, we measured contrast detection thresholds using the method of adjustment. The method of adjustment is not as accurate as a staircase procedure, but is sufficient for a quick check that all observers are able to perform the main experiment. To obtain detection thresholds, observers were asked to manipulate the contrast of a centrally presented grating using a trackball mouse. This grating shared the spatial parameters of the central grating used in the contrast matching procedure (shown in Figure 2). Rolling the trackball away from them increased the contrast, while rolling it towards them decreased the contrast in equal linear increments. Observers were given as much time as they needed to adjust the contrast of the grating until it was just barely visible, which they indicated by a keypress. While observers were adjusting the contrast, the grating slowly cycled on and off (0.15s ON : 2.15s OFF), and its orientation was randomised on every cycle. This procedure was repeated six times, with the median of these six thresholds taken as the observers absolute contrast detection threshold.

**Figure 2:**
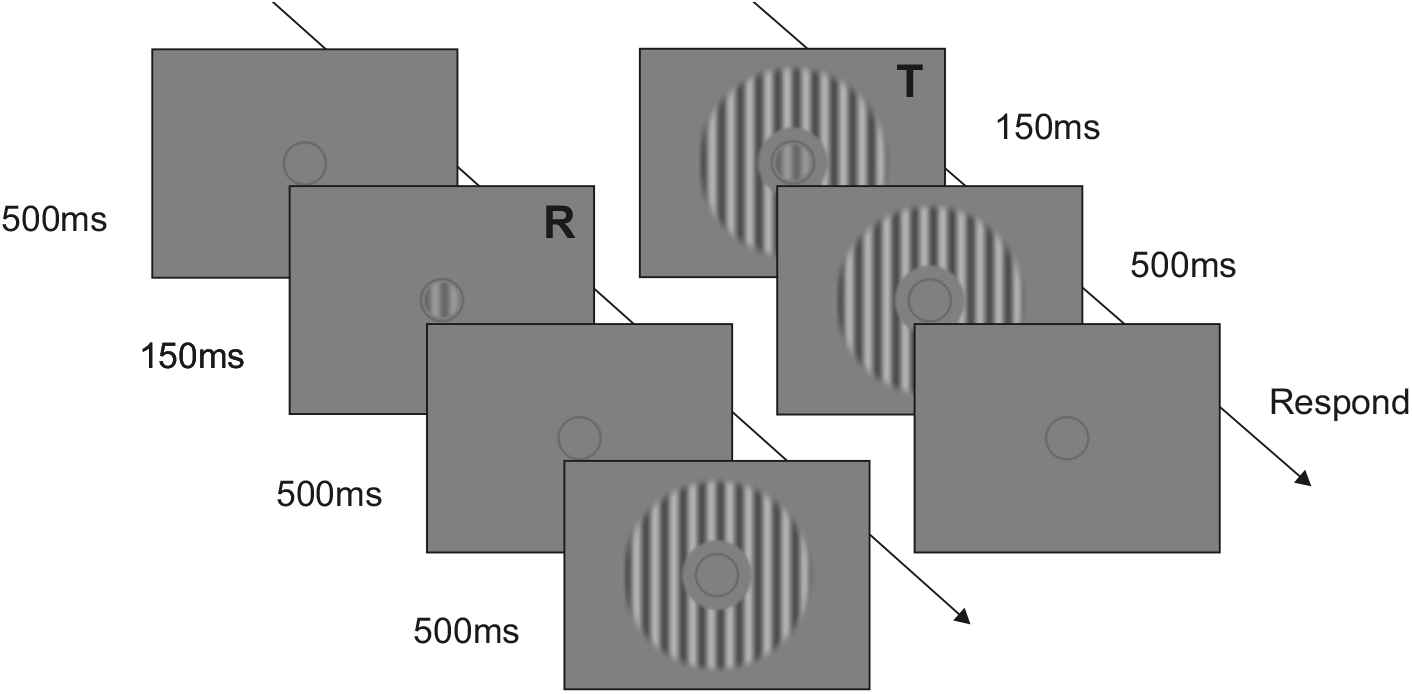
Timing of stimulus presentation. To be read from top-left to bottom-right, following diagonal arrows. The reference interval is represented by the “R” text, while the target interval is represented by the “T” text. These interval indicators are for illustrative purposes only and were not presented to the observer.

To assess the strength of surround suppression across different surround configurations, observers took part in a 1-up 1-down staircase procedure designed to measure the point of subjective equivalence (PSE) between the perceived contrast of an unmasked reference grating and a surround-masked target grating. Observers viewed target and reference gratings sequentially via a two-interval forced-choice (2IFC) procedure, and their task was to indicate the interval (1st or 2nd) that contained the central grating with the highest contrast. Many observers were completely unfamiliar with visual psychophysics, so we wanted to ensure that the experiment was as simple as possible. For this reason, the reference (unmasked) grating was always presented in the first interval (at 20% contrast), and the masked target grating was always presented in the second interval, with a contrast dictated by the staircase procedure over the course of 70 trials. Note, the correct response could be in either interval, as the contrast of the target grating can be greater or less than that of the reference grating. The target either had no mask (a control condition), a collinear mask, or an orthogonal mask, and surrounds were fixed at 40% contrast. The phase and orientation of presented stimuli was randomised such that the reference grating in interval one had a different phase and orientation to the grating and annulus presented in interval two. This was done to ensure observers were only responding based on the perceived contrast, and to prevent adaptation to the spatial parameters of the stimulus. As shown in Figure 2 a single trial consisted of the reference grating for 150ms, followed by an inter-stimulus-interval of 500ms. Observers then saw 500ms of surround, followed by 150ms of target grating and surround, and a final 500ms of surround alone. We included this 500ms onset delay for the target grating to prevent any contribution from temporal onset masking (Petrov & McKee, 2009). Observers were then given unlimited time to indicate (via a button-press) the interval that contained the central grating of highest contrast. The first 5 trials of the staircase procedure had a step-size of 2.25% contrast, but subsequent trials were reduced to a step-size of 0.75% to approximate the observer’s PSE more closely. Each of the surround-on conditions were repeated up to 3 times by each observer. If the PSE estimates of the 1st and 2nd run (based on a least-squares Weibull fit) differed by less than the minimum staircase step-size (0.75%), the final repetition was skipped. The no-surround control contrast matching condition was always conducted first, and only once. For each observer, this resulted in 5 – 7 completed staircases. To give observers the opportunity to obtain a stable response criterion prior to the main experiment, each practised with an abbreviated version of the experiment containing one surround and one no-surround condition, each with 25 trials of the staircase procedure. Additional practice runs were permitted if necessary. The entire experiment was completed in a single session in all cases, and each session took no more than 1.5 hours.

### Model fitting and statistical analysis

Observers’ final PSEs were estimated by fitting a logistic cumulative density function to accuracy estimates. This was accomplished using the maximum likelihood fitting routines of the Palamedes Toolbox (http://www.palamedestoolbox.org/) in MATLAB R2018b. The threshold and slope of this psychometric function were allowed to vary as free parameters, while the guess-rate and lapse-rate were both fixed at 0%, such that the threshold parameter equates the contrast achieving an accuracy of 50% (the PSE). For each observer, and each condition, this fitting procedure was carried out on the accumulated accuracies from all staircase repetitions by summing the number of correct responses and total trials. The effect of age group and surround configuration was investigated via a two-way mixed-model analysis of variance (ANOVA), with age group (younger, older) used as the between subjects factor, and surround condition (no surround, collinear, orthogonal) used as the within subjects factor. In younger observers, for the orthogonal condition, there was a violation of the assumption of normality of residuals (Shapiro-Wilk, p *<* .001) due to the presence of an outlier - how we dealt with this data-point is detailed in the Results section. This dataset satisfied Levene’s test for the homogeneity of variance (p *>* .05), but failed Mauchly’s test of sphericity (*χ*2 (2) = 29.03, p *<*.001), so the Greenhouse-Geisser correction was applied. To emphasises any within-subjects effects, we also supply an alternative illustration of PSEs as “suppression factors”, but will rely on the statistics carried out on the PSEs for inference. The suppression factor is simply the PSE for a given surround condition divided by the the PSE for the no surround condition. For completeness, we also report a comparison between younger and older observers’ contrast detection thresholds. Due to the presence of two outliers (beyond 1st and 3rd quartiles -/+ 1.5 * inter-quartile range), this comparison was made using an independent samples Mann-Whitney U test (comparing medians). All statistical analyses were carried out using SPSS version 28 (https://www.ibm.com/products/spss-statistics).

## Results

### Elevated contrast detection thresholds in older observers

First, to demonstrate that all observers were able to perceive the suprathreshold stimulus, we report their contrast detection thresholds. Absolute detection thresholds are shown in Figure 3, and are all well below the contrast of our matching stimulus (20%). However, there are two high-threshold outliers in the older age group, and a single marginal outlier in the younger age group. It is unclear whether these reflect genuine visual deficit, or whether they represent a misapprehension/artefact of the method of adjustment. Nevertheless, we have no reason to suspect that these data-points are artefacts, so we retained their thresholds and conducted a non-parametric comparison. We also retained these observers’ supra-threshold contrast matching data, as the contrast of the reference grating would still have been approximately twice their detection threshold. On average, younger observers had lower detection thresholds (median = 2.02%, IQR = 0.60%), than older observers (median = 3.55%, IQR = 1.74%), a difference in medians of 1.53% that was significant according to a Mann-Whitney U test (U = 28, z = -4.552, p *<* .001). This age-related increase in contrast detection thresholds is consistent with existing literature (Beard et al., 1994; Elliott, 1987; Mei & Leat, 2007; Owsley et al., 1983). Having confirmed that all observers could readily perceive our suprathreshold stimulus, we will now report the results of the contrast matching experiment.

**Figure 3:**
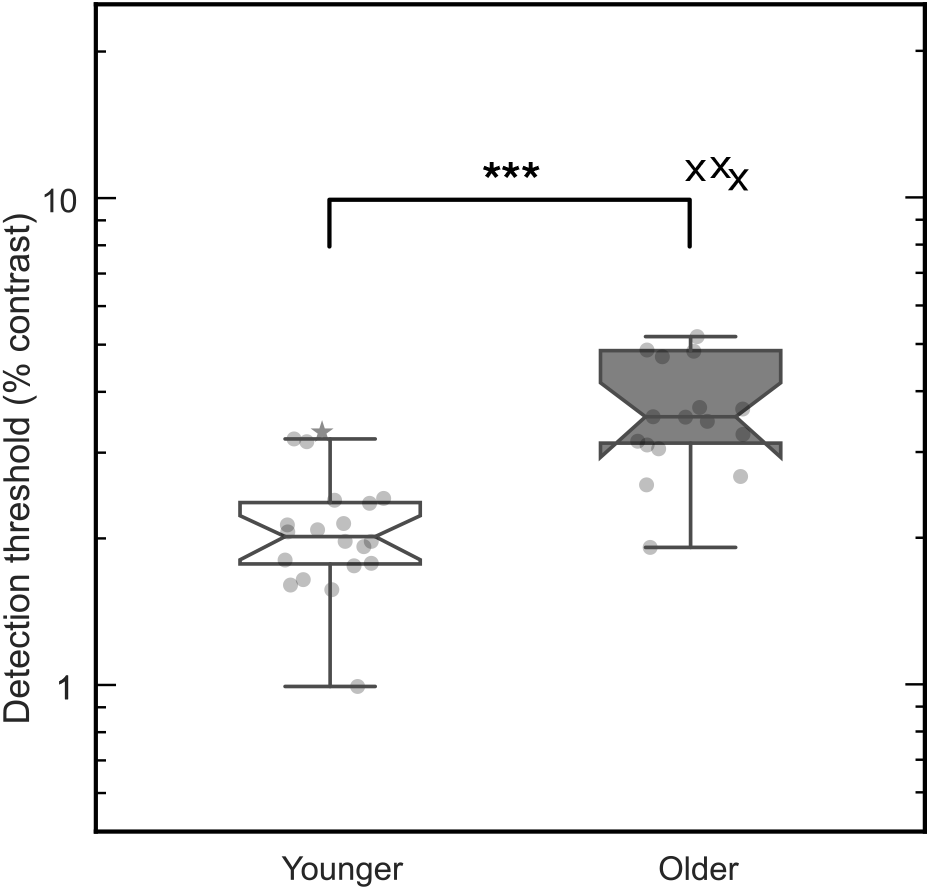
Contrast detection thresholds in younger and older adults represented as box-plots due to the use of non-parametric statistics to compare medians. Grey circles represent individual data points, crosses represent outliers. *** = p*<*.001. The upper whiskers of each boxplot extend to the last data-point less than the 3rd quartile + 1.5* the interquartile range (IQR), and the lower whiskers extend to the last data-point more than the 1st quartile – 1.5*IQR.

### Surround suppression does not change with age

In Figure 4, we present example psychometric function fits from a younger and an older observer. Both have discrimination accuracy estimates that are well described by a logistic cumulative density function, and the same was true of most observers. However, two observers had chance-level accuracy at high target contrasts that were not consistent with their performance at lower contrasts. In these observers, this discontinuity in performance was driven by several target intensities with only two trials, likely reflecting a brief excursion of the staircase into high target contrasts due to a series of response errors. For this reason, we removed any target intensities with fewer than three trials prior to descriptive model fitting for these two observers.

**Figure 4:**
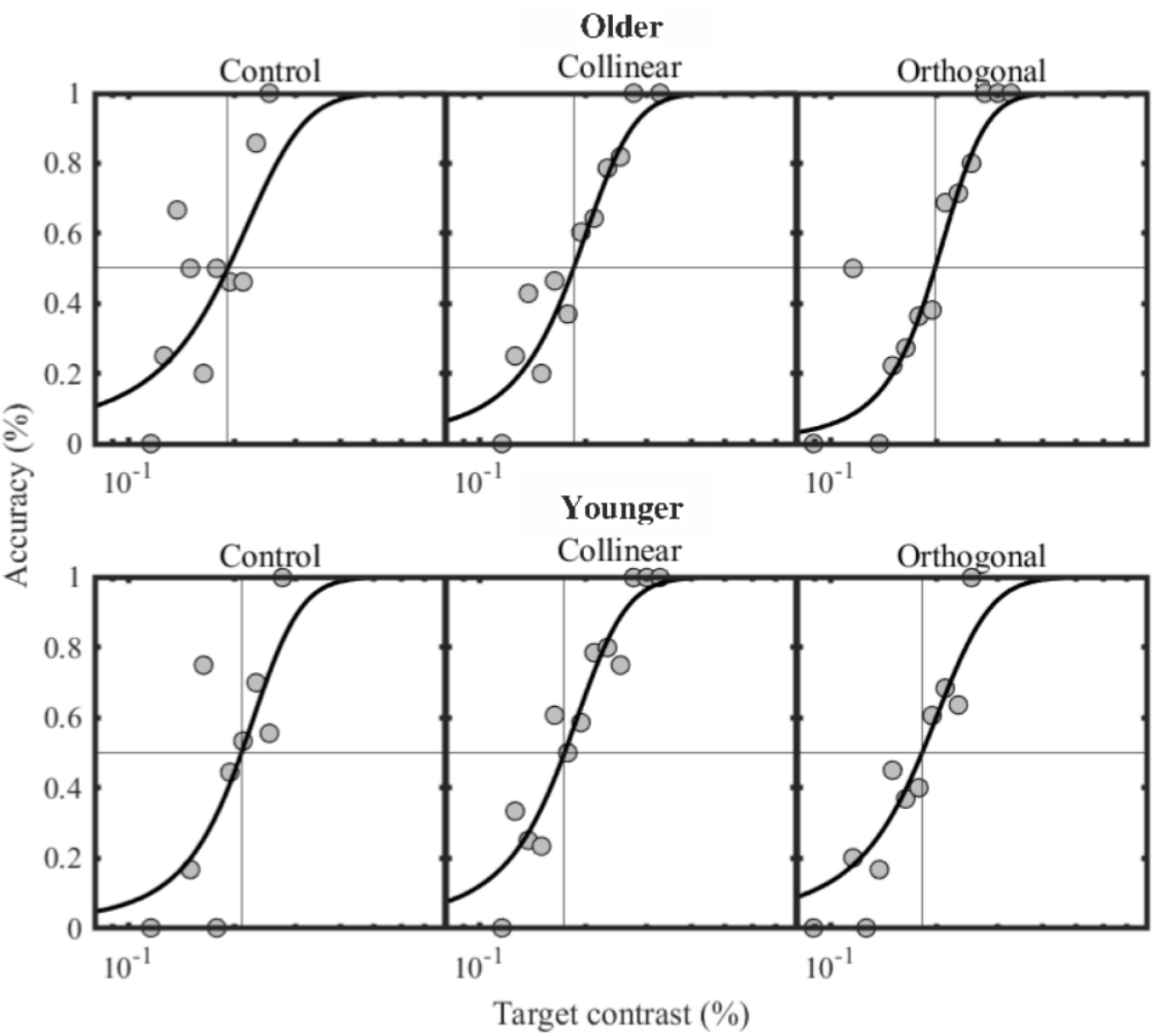
Psychometric function fits from two example observers in the three matching conditions we used. Where accuracy is at 50%, and where the vertical line of the cross-hair intercepts the abscissa denotes the point of subjective equality (PSE) that we use for group-level comparisons.

In Figure 5A, the mean of PSEs estimated by psychometric function fits for each surround condition are presented split by age group. We analysed observers’ PSEs with a mixed-model ANOVA, but examination of model residuals (via quantile-quantile plots) showed that there were two clear outliers (a studentised residual of ± 3) - one in the no-surround condition (older) and one in the orthogonal condition (younger). Further inspection revealed that the generative psychometric functions for these thresholds had sensible fits. As we had no reason to suspect these data-points were artifactual, we elected to run the ANOVA with and without their data to examine whether the statistical outcome was contingent on their contribution. The interpretation of all p-values was the same in either case, so we elected to interpret results based on the full dataset. In 5A, older and younger observers appear to have similar PSEs in all surround conditions, though there is some indication that younger observers’ PSEs may be more sensitive to the orientation of the surround. Statistical analysis showed that the within-subjects comparison of surround condition was significant (F(1.280,42.240) = 26.545, p *<* = .001), while the between subjects comparison of age group was non-significant (F(1,33) = .060, p = .808), and there was no significant interaction between age group and surround condition (F(1.280,47.240) = .273, p = .662). These results indicate that PSEs were contingent on the configuration and/or presence of the surround, but do not support the presence of a broad age-related effect on surround suppression. Moreover, our results do not support any surround-configuration-specific effect of age. This pattern of results is better conveyed in Figure 5B, where we have plotted surround-on PSEs as multiples of observers’ surround-off PSEs, emphasising within-subjects differences relative to the surround-off control. Here, we can see that 95% confidence intervals on the mean do not cross 1 for both surround configurations for both age groups (indicative of a significant PSE reduction from the surround-off control), but are almost entirely overlapping within each surround condition, implying a similar degree of surround suppression across age groups.

**Figure 5:**
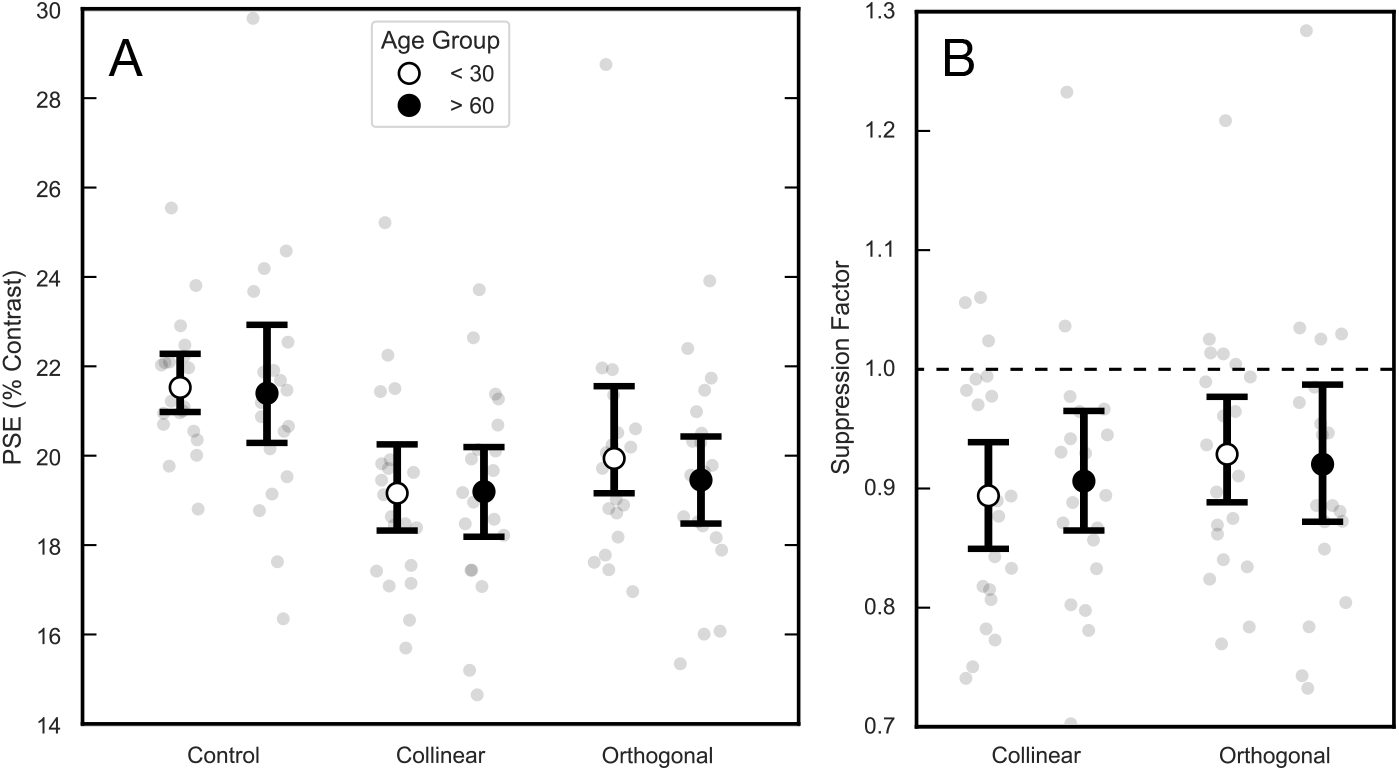
A: mean of PSEs split by age group and surround condition. Larger open circles represent the within condition mean of young participants, larger closed circles represent the same for older participants. Error bars represent 95% confidence intervals. B: Same format as A, but showing suppression ratios relative to individual contrast matches in the absence of a surround. Thresholds below the solid reference line are indicative of suppression.

As the within-subject effect of surround condition was significant, we conducted post-hoc paired t-tests across all observers, collapsed across age groups (Figure 6A). Observers’ PSEs were, on average, highest in the no-mask control condition (mean = 21.5%, 95%CI =[20.8, 22.3]), lowest in the collinear mask condition (mean = 19.2%, 95% CI [18.6, 19.9]) and intermediate in the orthogonal mask condition (mean = 19.7%; 95CI = [19.1, 20.6]). The mean reduction in PSEs between the control and collinear conditions was significant (mean Δ = -2.3%, 95% CI: [1%, 3%] p *<* .001), as was the reduction in PSEs between control and orthogonal conditions (mean Δ = -1.8%, 95% CI [.09%, 2.6%] p *<*.001), and the difference between orthogonal and collinear surround PSEs (mean Δ = 0.5%, [.03%, 1.3%], p *<* .05), though the latter was very small - less than the smallest step-size of our staircase procedure. As before, these PSEs are replotted as suppression factors in Figure 6B, where, on average, collinear mask PSEs were 90% (95% CI = [86, 93]) of the no mask control, increasing to 92% (95% CI = [89, 96]) with an orthogonal mask, representing a very mild reduction in masking strength for orthogonal surrounds.

**Figure 6:**
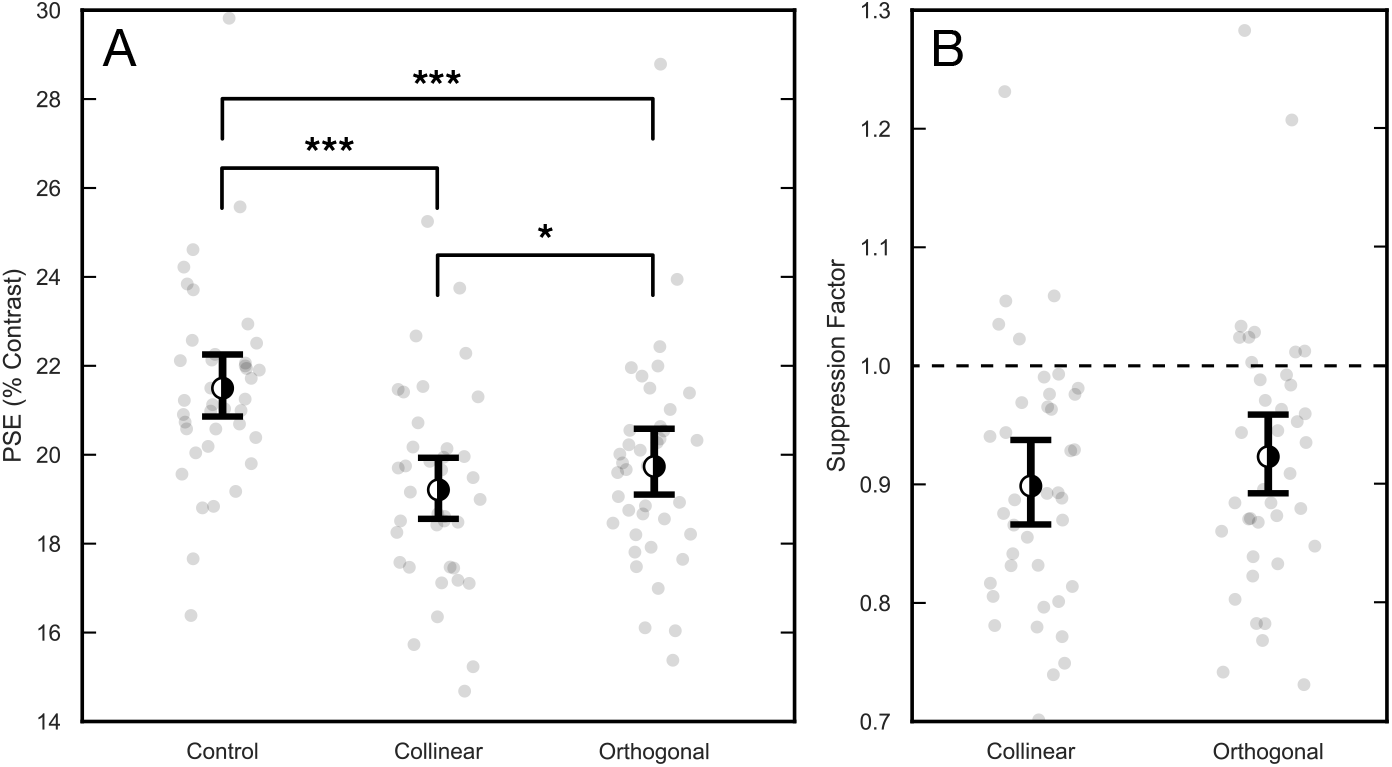
A: mean of PSEs collapsed across age groups, split by surround condition. Half-filled circles represent the within-condition mean across all observers. Smaller grey dots represent individual data points. Error bars represent 95% CIs * = p*<*.05, *** = p*<*.001. B: mean of suppression factors, collapsed across age groups. As before, points below the horizontal reference line represent surround suppression.

## Discussion

Here, we asked whether age alters a fundamental component of early visual processing in humans: surround suppression. Following previous reports, we used foveally presented stimuli, but modified them to minimise edge effects and overlay masking. We found significant surround suppression in both older and younger age groups. However, the suppressive effect and spatial tuning was weak compared to the effects found in the periphery both at detection threshold and suprathreshold (Petrov et al., 2005; Shushruth et al., 2013; Xing & Heeger, 2000). Crucially, we found no difference in the strength of surround suppression between younger and older participants. We will now discuss where our results lie in the context of the previous literature on surround suppression and offer alternative explanations for the age dependence that has been previously reported.

### Untuned foveal surround suppression

Previous work has reported foveal surround suppression at supra-threshold contrasts that is orientation tuned (Cannon & Fullenkamp, 1991; Nguyen et al., 2020; Xing & Heeger, 2000, 2001). We too found significant supra-threshold suppression, but found a smaller (though significant) difference between collinear and orthogonal suppression indices. What might explain the weaker tuning we have encountered? One explanation for our differing observations is the spatial parameters of our respective stimuli. Our stimulus, subtending 3.2 degrees of visual angle, is still smaller than the central probe used in both of Xing and Heeger’s experiments (which subtended 3.5 degrees of visual angle). Additionally, their surrounds subtended 11° (Xing & Heeger, 2000) and 7° (Xing & Heeger, 2001), while ours only subtended 3.2°. So it is plausible that their stimuli recruited longer-range suppression that may be distinct in its tuning characteristics. Furthermore, Cannon and Fullenkamp (1991) and Nguyen et al. (2020) both used stimuli similar to ours (in both spatial configuration and contrast) and both report that orthogonal surrounds reduced the strength of suppression by a factor of approximately 1.5, whereas we report a minute reduction in strength by a factor 1.03. The only consistent difference between our work and existing similar work is the inclusion of a one-grating-cycle gap between the center and the surround: all other reports mentioned used closely abutting surrounds. Thus, some contribution from overlay masking cannot be discounted in previous work. Additionally, while overlay masking is not as strongly orientation tuned as surround suppression in the periphery, it is still broadly tuned (Petrov et al., 2005). This means the mild orientation tuning (mild relative to the periphery) found previously could just be a manifestation of broadly tuned overlay masking at the fovea. Future work investigating the orientation tuning of overlay masking at the fovea may explain our discrepant results.

We turn now to the temporal parameters of the stimuli used. Xing and Heeger (2000, 2001) used central probe gratings that contrast reversed at 8Hz and were presented for 500ms in the context of static surrounds. (Cannon & Fullenkamp, 1991) used static central gratings and surrounds, and presented them for 1 second. Like Nguyen et al. (2020), our central probe stimuli were presented for 150ms, but our surrounds appeared 500ms in advance of the probe and persisted for 500ms after the probe. Thus, we used shorter duration central probe stimuli than most previous reports, apart from Nguyen et al. This may mean that the contrast of our probes were suppressed by a different (more transient) mechanism. In the periphery, both transient and sustained surround suppression are found to be orientation tuned (Petrov & McKee, 2009), and the suppression imposed on briefly presented stimuli (less than 100ms) is strongest. It could be that the temporal invariance of orientation tuning Petrov and colleagues report in the periphery does not persist at fixation. However, that Nguyen et al. (2020) still find moderate orientation tuning is contrary to this hypothesis, as they too used short-lived stimuli. A remaining possibility is that onset masking is orientation tuned, and that the 500ms early onset of the surround that we used escaped this form of suppression, leaving only the suppressive input from a weaker less-tuned mechanism. At 8 degrees eccentricity Petrov and McKee (2009) measured the effect of presenting the surround in advance of the target, and found that the strength of surround suppression rapidly decayed when the surround is presented as little as 20ms in advance of the target, but is still orientation tuned. However, in their early onset surround conditions, the surround was removed before the target was presented, so their results do not indicate whether the early-onset but persistent surround we used would have negated orientation-tuned suppression, which might be expected if the inhibitory pool is driven mostly by neurons with transient receptive field properties.

### Alternative explanations for age effects in contrast suppression

Previous work has indicated that surround suppression of contrast at suprathreshold intensities increases with age (Karas & McKendrick, 2009, 2011, 2012). We did not replicate this finding, and propose that the age-related alteration of a different suppressive mechanisms could explain our discrepant results. The idea that surround suppression is not the only explanation is compatible with several of the findings in previous reports. First, more recent work has found that the age-related increase in central-field suppression is abolished by inter-ocular presentation of the centre and surround (Pitchaimuthu et al., 2017), indicating that the inhibitory circuitry affected by age exists prior to the fusion of the monocular visual streams in V1. The authors suggest that the mechanism affected by age could be a separate, ‘early’ surround-suppression component, such as that proposed by Webb et al. (2005) (based on macaque V1 electrophysiology) which is more apparent at low contrasts 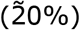, where the age effect has been shown to be strongest in humans (Karas & McKendrick, 2015). However, the same pattern of results could also be explained by overlay masking. It too has a broadly tuned monocular component (Baker et al., 2007; Bonds, 1989; DeAngelis et al., 1992), and has been shown to occur prior to orientation-tuned surround suppression (Petrov et al., 2005). It is important to note that these previous findings for overlay suppression are at detection threshold, and it is uncertain how far they extend to suprathreshold contrast perception.

It is also possible that a broadly orientation tuned overlay masking contributed to previous findings.Nguyen and McKendrick (2016) demonstrated that a very narrow surrounding annulus presented in close proximity to a central grating (a gap of *∼λ*/3) is sufficient to produce increased suppression in the elderly – approximately 70% of the age-effect they reported when using spatially extensive directly abutting surrounds at the same contrasts in another experiment (Karas & McKendrick, 2015). This implies that the wide-field extent of the surround is of less importance relative to the inner boundary, and indeed, in the same work they found the inverse age-effect in the periphery when they used a intervening gap in excess of 1 stimulus cycle. While this was interpreted as a foveal bias in their observed age effect, any contribution from overlay suppression would have been reduced in their peripheral condition. However, recent work did not find the age effect even when using directly abutting surrounds (Nguyen et al., 2020), suggesting that a general contribution from overlay masking is not be the sole explanation for the differences in our findings. In this experiment, the authors used stimuli at lower spatial frequencies than usual (1cpd), where a sizeable proportion of the surrounding annulus would be within one stimulus wavelength of the central target. That the effect of age was not found in this case suggests that there may be some selectivity in the mechanism affected by age, such that it is absent at low spatial frequencies. If we accept that surround suppression changes in spatial tuning as a function of absolute stimulus contrast (Levitt & Lund, 1997; Webb et al., 2005), then different suppressive mechanisms may be dominant at low spatial frequencies which may escape the effect of age.

Another explanation for the absence of an age-related strengthening of surround suppression in our results could be the timing of our stimuli. Our central targets were presented for 150ms, a duration shown to produce the age-effect previously (Nguyen & McKendrick, 2016). As mentioned previously, our stimuli are unique in that the onset of the surrounding annulus was not simultaneous with the onset of the central target. Previous magnetoencephalography (Ohtani et al., 2002) and electroencephalography (Schallmo et al., 2018) experiments, have presented surrounds up to 2s before and after the target (to measure the separate responses to the centre and surround), and still measure significant suppression of neuronal responses. In a control condition, Ohtani and colleagues even compared the suppression elicited by simultaneous and sustained early-onset surrounds, finding similar suppression in both cases. Moreover, in a psychophysical experiment, Schallmo and Murray (2016) presented surrounds 1-2s in advance of a probe until 500ms after the probe’s offset, and still reported surround suppression at suprathreshold contrasts. Schallmo and colleagues’ stimuli were presented in the perifovea (5.3/°), as were those of Ohtani et al., so there remains the possibility that this pattern of results may change at fixation. Nevertheless, if the effect of age on surround suppression in central vision requires the simultaneous onset and/or offset of both the centre and surround, our stimuli would have escaped it. As stronger surround suppression with age has been identified using centres and surround presented for as long as 500ms (Karas & McKendrick, 2015), we think this is unlikely, but investigating the temporal dependencies of this effect would be an interesting line for enquiry. Some temporal specificity is consistent with previous reports showing that the increase in suppression with age is present for intermediate durations of 150 – 200ms but severely diminished at longer durations of 500ms and totally absent at shorter durations of 40ms (Karas & McKendrick, 2015; Pitchaimuthu et al., 2017). Indeed, that such minor alterations to the stimulus (including our own) abolish the age effect suggests that it is generated by a suppressive mechanism (or a subset of mechanisms) that is spatially and temporally narrow-band, rather than a general feature of contrast gain-control processes throughout the visual system.

## Conclusion

We have shown that the age-related increase in surround suppression reported in a recent series of publications is not a universal finding. Using stimuli designed to measure surround suppression we report no effect of age on supra-threshold contrast suppression, despite using contrasts previously reported to yield the strongest age-related difference. We propose that the discrepancy between our results and previous reports could be the narrow-band spatiotemporal tuning of the mechanism effected age or unintentional contributions from overlay masking. The suppression we did find is weak, and only weakly tuned to the collinearity of the target and the surround, unlike previous reports of suprathreshold masking in central vision.

## Notes

### Competing Interest Statement

The authors have declared no competing interest.

